# Protein secondary structure affects glycan clustering in native mass spectrometry

**DOI:** 10.1101/2021.05.01.442239

**Authors:** Hao Yan, Julia Lockhauserbäumer, Gergo Peter Szekeres, Alvaro Mallagaray, Robert Creutznacher, Stefan Taube, Thomas Peters, Kevin Pagel, Charlotte Uetrecht

## Abstract

Infection with human noroviruses (hNoV) for the vast majority of strains requires attachment of the viral capsid to histo blood group antigens (HBGA). The HBGA binding pocket is formed by dimers of the protruding domain (P dimers) of the capsid protein VP1. Several studies have focused on HBGA binding to P dimers, reporting binding affinities and stoichiometries. However, nuclear magnetic resonance spectroscopy (NMR) and native mass spectrometry (MS) analyses yielded incongruent dissociation constants (K_D_) for binding of HBGAs to P dimers and, in some cases, disagreed whether glycans bind at all. We hypothesized that glycan clustering during electrospray ionization in native MS critically depends on the physicochemical properties of the protein studied. It follows that the choice of the reference protein is crucial. We analyzed carbohydrate clustering using various P dimers and eight non-glycan binding proteins serving as possible references. Data from native and ion mobility MS indicate that the mass fraction of β-sheet has a strong influence on the degree of glycan clustering. Therefore, the determination of specific glycan binding affinities from native MS must be interpreted cautiously.

## 1. Introduction

Infection with human norovirus (hNoV) is the most common cause of acute gastroenteritis, leading to an estimated 685 million cases annually worldwide, with elderly, immunocompromised patients and children under 5 years most severely affected. hNoV belongs to the family of *Caliciviridae*, non-enveloped viruses of icosahedral shape with the viral genome consisting of positive-sense single-stranded RNA. Histo blood group antigens (HBGA) serve as attachment factors in viral infection [1,2]. Previous NMR, X-ray crystal structure and mass spectrometry (MS) investigations identified the L-fucose moieties within HBGAs as minimal binding motif for hNoV attachment. Notably, L-galactose derived from L-fucose by substitution of one hydrogen atom with a hydroxyl group at C6 is known not to bind [3,4]. As determination of dissociation constants and binding stoichiometries is key in understanding protein-ligand interactions a number of studies reported such data for binding of HBGAs to hNoVs based on different experimental approaches. Sun *et al*. previously established an MS-based method to determine specific glycan binding to proteins [5]. The method relies on the addition of reference proteins into the sample mixture and simultaneous analysis using native MS. This approach allows the quantitation and elimination of unspecific ligand clustering during the electrospray ionization (ESI) process. Clustering arises from statistical presence of free ligands within the same droplet as the free or ligand-bound proteins, which can then dry down to the protein surface upon droplet evaporation. This process becomes relevant at elevated ligand concentrations and is independent of protein molecular weight [6-8]. Calculations to correct for this effect are based on total peak areas per mass species within the same spectrum to ensure identical ionization conditions.

A couple of groups including us have employed this method to characterize glycan binding to hNoV P dimers using distinct reference proteins [4,9-12]. Notably, results disagreed [4,11], raising the question whether selection of the reference protein influences data interpretation. Moreover, with the exception of glycan mimetics [13], the search for non-binding ligand or P dimer controls for native MS was unsuccessful in our hands. This is in stark contrast to NMR data, which shows that certain P dimers and certain glycans do not interact. This suggests a severe issue with the chosen native MS approach. Strikingly, direct MS analysis had determined mM K_D_s for several sialic acid containing carbohydrates binding to hNoV P dimers [9,11,12,14], whereas orthogonal STD and protein based CSP NMR experiments clearly revealed no binding of sialic acids to hNoV P dimers and virus-like particles (VLPs) [9,15,16]. This questions the validity of the results from direct MS measurements.

Additionally, the reported K_D_ values from carbohydrate binding studies to P dimers are not comparable in STD NMR, native MS and isothermal titration calorimetry (ITC) [11,12,17]. Importantly, K_D_s obtained for active pharmaceutical agents based on different biophysical assays such as native MS, ITC, surface plasmon resonance (SPR) and circular dichroism (CD) were equivalent in numerous other cases [8,18,19]. However, these compounds showed higher binding affinities (µM range) towards the target protein and exhibited in most cases no carbohydrate-like structures [20]. In contrast, glycan binding affinity is often in the high µM to mM range. This also holds for hNoV carbohydrate interactions [9-12,14,21]. In former studies, the origin of the discrepancies between NMR and native MS data for P dimer-glycan interaction was not in focus [12]. The problem became evident when comparing wildtype, i.e. non-deamidated, and deamidated GII.4 Saga P dimers. In this GII.4 P dimer, an asparagine residue flanking the binding site is specifcally and spontanously converted into an iso-aspartate. The deamidated P dimer has been shown to have greatly reduced glycan binding affinity in NMR and hydrogen/deuterium exchange (HDX) MS, roughly by an order of magnitude for HBGA B trisaccharide and fucose compared to the wildtype [4,13].

Here, wildtype and deamidated P dimers are compared using native MS to shed light on potential methodological issues. Moreover, Gb4, an all-galactose tetrasaccharide, is employed as negative control. Using this information, we hypothesize that clustering depends on physicochemical properties of the proteins. To deduce what obscures the binding studies, multiple reference proteins varying in their properties are compared. The results point to an influence of β-sheet content. This theory is corroborated by ion mobility MS (IMMS) measurements on P dimers in presence of glycans and additional data on other P dimers, *e*.*g* from MNV P dimers, which were recently shown by NMR not to bind glycans at all [22]. Our results indicate that reference proteins need to be chosen carefully to match structural properties of the target protein for glycan binding studies and crucially, suggest an additional influence of structural dynamics, which preclude glycan binding studies in native MS for hNoVs.

## 2. Materials and Methods

### 2.1 Glycans

The following glycans were purchased from Elicityl-Oligotech, dissolved in H_2_O for native MS analysis and are shown in Fig. 1. Blood group antigens: **(1)** A tetrasaccharide type 1 (>90% NMR) (GalNAcα1-3(Fucα-2)Galβ1-3GlcNAc, product code: GLY035-1-90%), **(2)** B tetrasaccharide type 1 (>90% NMR) (Galα1-3(Fucα1-2)Galβ1-3GlcNAc, Product code: GLY038-1-90%). Globo-series: **(3)** P antigen Gb4 (>90% NMR) (GalNAc β1-3 Galα1-4Galβ1-4Glc, Product code: GLY121-90%). The HBGA ligands were chosen as known binders based on their fucose binding moiety, whereas Gb4 is the fucose-free non-glycan-binding reference. HBGAs comprise various oligosaccharides, the tetrasaccharides were chosen here instead of e.g. trisaccharides to have a larger mass increment upon association with the P dimer allowing for more gentle conditions in the native MS measurements.

**Figure 1.**
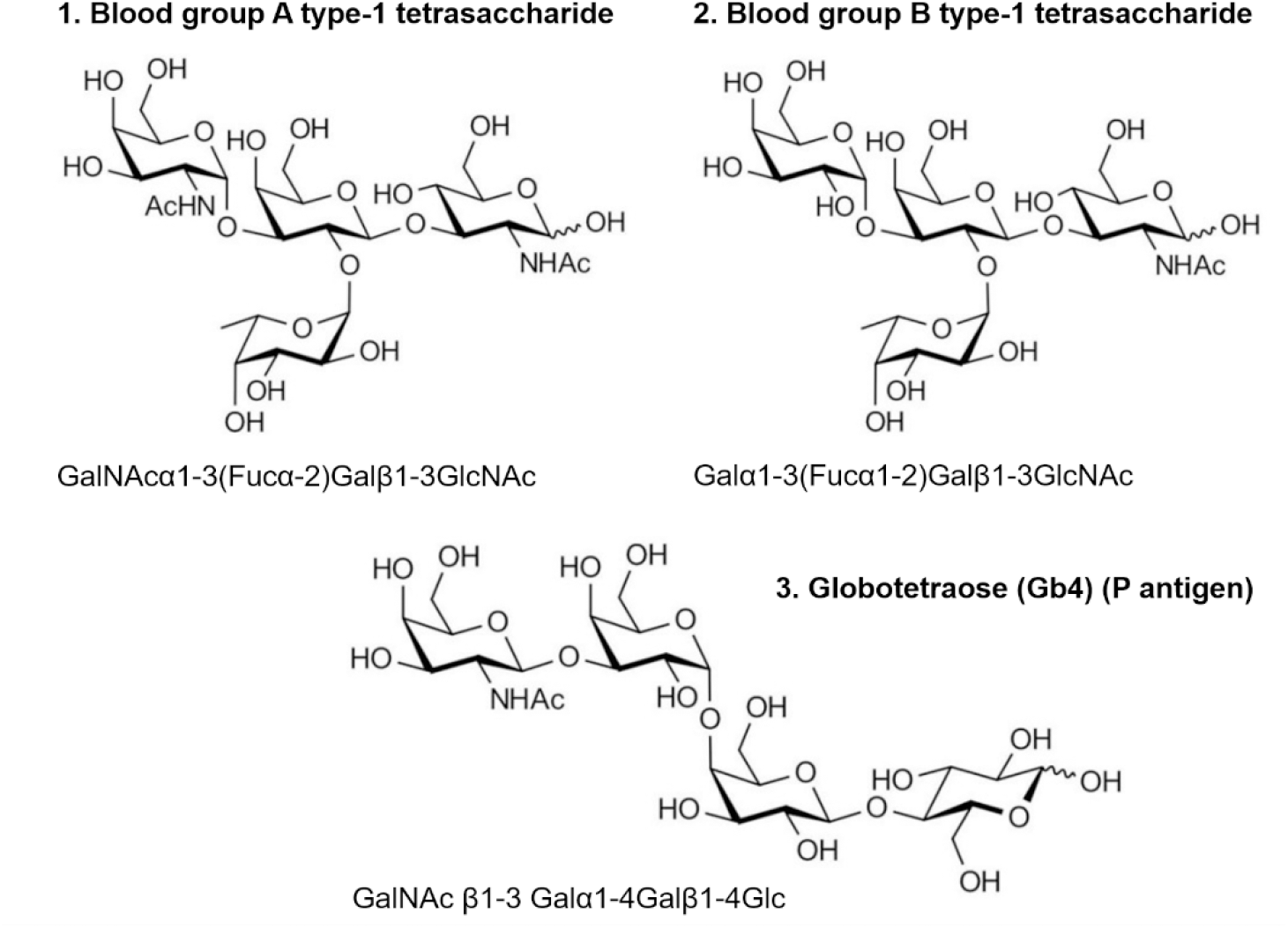
Structure of used carbohydrates. Blood group A and B type-1 tetrasaccharides (HBGAs) are known to bind to hNoV P dimers, Gb4 is considered a non-binder.

### 2.2 Proteins

P dimers of hNoV VP1 were used as target proteins. Native MS investigations were performed with P dimers from the hNoV strains GII.4 Saga (Saga 2006 (GenBank ID: AB447457, aa 225–530), GII.4 MI001 (KC631814, aa 225–530) and murine noroviruses (MNV-1, CW1: DQ285629, aa 228–530). *Escherichia coli* was used for overexpression of the P domains as described in [4,22]. Purified P dimers were stored at 4°C before preparation for native MS experiments. The commercially available reference candidates cytochrome c from equine heart (cyt c, CAS number 9007-43-6), ubiquitin from bovine erythrocytes (Ubq, CAS number 75986-22-4), carbonic anhydrase isozyme II from bovine erythrocytes (CA, CAS number 9001-03-0), alcohol dehydrogenase from *Saccharomyces cerevisiae* (ADH, CAS number 9031-72-5), L-Lactic dehydrogenase from rabbit muscle (LDH, CAS number 9001-60-9), myoglobin from equine heart (Myo, CAS number 100684-32-0), human apo-transferrin (apo-TFF, CAS number 11096-37-0) were purchased from Sigma-Aldrich (Merck, Germany) and stored according to manufacturer recommendation. Furthermore, a superfolder green fluorescent protein (GFP) was kindly provided by Henning Tidow (University of Hamburg, Germany) and used as reference protein candidate [23].

### 2.3 Native mass spectrometry

Prior to MS analysis, purified P dimers were buffer exchanged to 150 mM ammonium acetate (Sigma-Aldrich, reagent grade ≥ 98%) at pH 7 via centrifugal filter units at 12000 *g* at 4°C (Vivaspin 500, MWCO 10000, Sartorius). Glycans for native MS analysis were mixed with 1 µM (per monomer) purified GII.4 Saga P dimer and 3 µM of the reference proteins at indicated concentrations. Native mass spectra were measured at room temperature in positive ion mode on a high mass modified LCT Premier mass spectrometer (Waters/Micromass, UK and MS Vision, the Netherlands) with a nano-ESI source [24]. In house made gold-coated electrospray capillaries were used for direct sample infusion without any accessory chromatographic separation. The voltages and pressures were optimized for non-covalent protein complexes. The gas pressure used in the source hexapole was set between 6.5 and 6.8 × 10^−2^ mbar argon optimized for minimal complex dissociation and the backing pressure of the source roughing pump was set between 7.0 to 8.5 mbar. Spectra were recorded with applied voltages for the capillary and cone of 1.20 kV and 240 V, respectively. For calibration of the raw data, a 25 mg/ml cesium iodide spectrum from the same day was used. MassLynx V4.1 (Waters) was used to assign peak series to protein species and to determine the mass after minimal smoothing. Furthermore, OriginPro 2016 SR2 (OriginLab) was used for spectral deconvolution and curve fitting. Correction for nonspecific protein-ligand clustering was performed as described [5]. Corrected peak areas were averaged and normalized to the free protein signal. Data are based on at least three independent measurements. Python code Panda and Numpy package were used to calculate the specific binding of P dimers according to the glycan clustering ratio obtained from the selected reference protein candidates. Seaborn and matplotlib.pyplot package were imported into a python script to plot the results. The module of sklearn.linear model was applied for linear regression. The matplotlib and heatmap package were used to plot the heat-map.

### 2.4 Determination of glycan binding: calculation of the dissociation constant (K_D_)

Data from native ESI MS measurements were translated into dissociation constants (K_D_ values). The detailed calculations are based on the equations listed below. A protein (P) and a glycan ligand (L) form a complex through non-covalent interactions leading to a reversible association described by (equation 1):

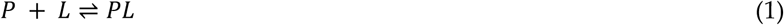

When the equilibrium is reached, the K_D_ can be calculated directly from the concentrations via the law of mass action (equation 2):

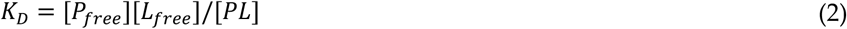

Theoretically, the K_D_ is calculated by measuring the concentration of the three components P_free_, L_free_, PL in solution. Usually, these values are only indirectly accessible requiring ligand titration up to binding saturation. However, native MS allows to directly measure P_free_ and PL, from which L_free_ can be calculated using the input concentrations. In the present work, the non-covalent interaction was measured in the gas phase assuming identical ESI and detection efficiency for free and glycan associated proteins, both for the analyte and the reference protein. The peak-areas (*A*) of the non-bound protein ion (*P*^*n+*^) and the bound protein ion (*PL*^*n+*^) were used to calculate the ratio (*R*) as these most accurately reflect the total signal and hence are assumed to be comparable to the ratio of a ligand bound protein complex and an unbound protein in solution (e.g. equilibrium state). For P dimers, the equations reflect glycan binding to the individual monomers [4] assuming that binding sites on either monomer are equal and independent [25].

Using *R* described in equation 3, the K_D_ calculation can be re-written into equation 4 with the known initial concentration of glycan ligand [L_0_] and protein [P_0_], adapted from Sun *et al* [5]:

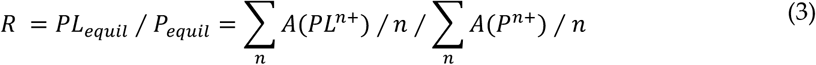

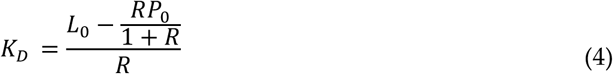

### 2.5 Titration measurement of glycan binding on P dimers

For the carbohydrate binding experiments, it is essential to keep the initial concentration of proteins and the pH value constant during analysis of protein-carbohydrate solutions. Therefore, accumulated spectra were obtained within 10 min after the voltages were applied to minimize the influence of pH changes during the nano-ESI process. The LCT mass spectrometer with only a short hexapole prior to the ToF analyser reduces potential activation and hence in-source dissociation (ISD). Three different glycan ligands including ligands of interest (HBGA B and A) and non-binding control (Gb4) were measured in a range of concentrations to determine the K_D_ of the first binding event of carbohydrates to P dimers (Tab. 1).

**Table 1.** Glycan clustering ratio *R* and K_D_ of HBGA B interaction with P dimers analysed by native MS. Data correction is based on the reference protein method.

| Ref. Protein | Mass /kDa | PDB ID | Secondary structure composition | | glycan clustering ratio $R$ at 500 $\mu M$ | | | $K_D$ for HBGA B /mM |
| --- | --- | --- | --- | --- | --- | --- | --- | --- |
| | | | % β-sheet | % α-helix | HBGA B | Gb4 | HBGA A | |
| Ubq | 8.9 | 1UBQ | 24.7 | 20.8 | 0.25±0.00 | 0.09±0.03 | 0.10±0.05 | 0.20±0.13 |
| Cyt c | 13.2 | 2N3B | 0 | 53.8 | 0.35±0.03 | 0.45±0.02 | 0.41±0.09 | 0.16±0.05 |
| Myo | 17.0 | 1AZI | 0 | 79.2 | 0.36±0.05 | 0.37±0.06 | 0.34±0.03 | 0.25±0.10 |
| GFP | 26.8 | 1BFP | 55.0 | 18.5 | 0.67±0.02 | 0.80±0.03 | 0.86±0.04 | 0.90±0.35 |
| CA | 29.1 | 1V9E | 35.8 | 15.8 | 0.75±0.07 | 0.75±0.03 | 0.74±0.07 | 0.40±0.10 |
| apo-TFF | 79.6 | 2HAU | 15.7 | 34.5 | 0.82±0.03 | 0.80±0.05 | 0.79±0.03 | n.a. |
| LDH | 146.8 | 3H3F | 17.1 | 46.9 | 1.02±0.04 | 0.91±0.06 | n.a. | n.a. |
| ADH | 147.0 | 4W6Z | 34.8 | 28.4 | 1.45±0.11 | 1.26±0.03 | 1.32±0.10 | 24.00±16.00 |
| SAGA P dimers | 68.1 | 4X7C | 32.5 | 6.6 | 1.55±0.05 | 1.40±0.21 | 1.44±0.15 | n.a. |

### 2.6 Ion mobility mass spectrometry (IMMS)

IMMS experiments on the GII.4 Saga P dimers were performed in a Synapt G2-S instrument modified with a linear drift tube ion mobility cell operating with He drift gas under 2.4 mbar pressure. Samples with 3 µM P dimer protein and 0, 100, 200, and 500 µM of the HBGA B tetrasaccharide or Gb4, respectively, were prepared, thoroughly homogenized, then loaded onto a Pd/Pt-coated borosilicate glass capillary. The sample was introduced by nano-electrospray ionization with the following instrumental parameters: capillary voltage – 1200 V; cone voltage – 15 V; source offset – 15 V; source gas flow: 0.0 mL/min; backing pressure – 0.1 bar; source temperature – 30 °C; these settings ensured the soft ionization conditions that allow for preserving the native protein structure. The arrival time distribution curves shown in Figure 5 were recorded at a He cell DC of 35 V and a bias of 55 V. Each experiment was performed at least twice to address reproducibility.

## 3. Results

### 3.1 Glycan clustering on norovirus P dimers

Here, the wildtype and deamidated GII.4 Saga P dimers are used as positive and negative protein binding control [4], respectively. The tetrasaccharide of HBGA B type 1 is employed as glycan known to bind to the wildtype and Gb4 as an all galactose glycan non binder [9,10]. Thereby, the applicability of the MS approach to investigate binding equilibria in presence of a reference protein is verified. As can be seen in Fig. 1A, the reference protein cytochrome c (cyt c) picked up similar amounts of clustered glycans in both spectra indicating similar spray conditions. Surprisingly, also non-deamidated and deamidated P dimers revealed a comparable pattern of glycan attachment, although a reduced amount of glycans was expected to stick to the deamidated protein. After correction for clustering to the reference protein, even more glycans are supposedly specifically bound to the deamidated P dimer, which serves as low/non-binding control, than to the non-deamidated. Moreover, occupancy is much higher than what would be expected based on the K_D_s for the wildtype determined by NMR (12 mM for B tetrasaccharide [22]) suggesting an intrinsic issue with the measurement approach. Additionally, the non-binding glycan Gb4 showed a pattern similar to HBGA B after correction (Fig. 1B), however, with more glycan attached to the wildtype protein.

**Figure 1.**
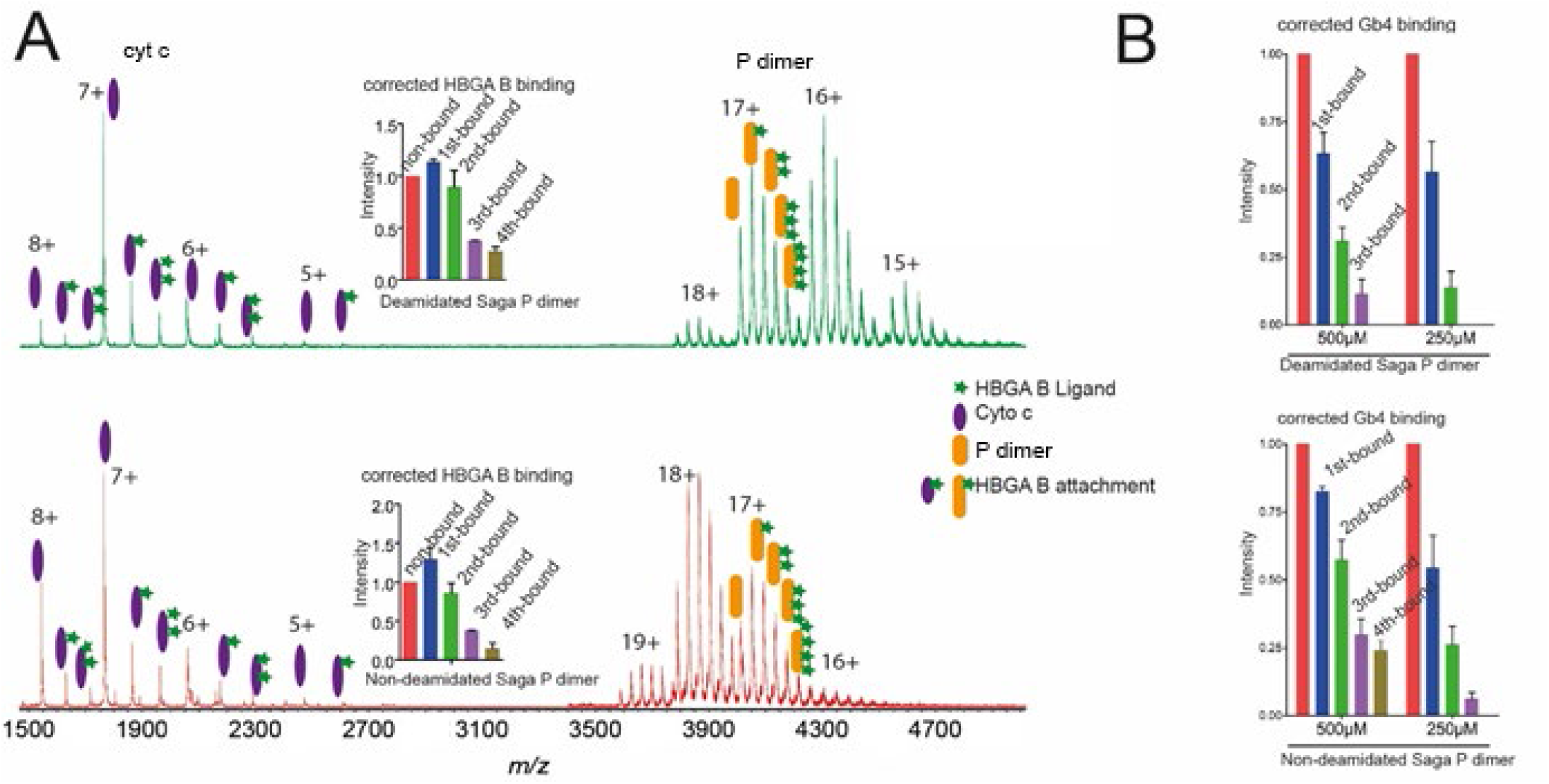
Influence of deamidation on glycan clustering. (A) Native mass spectra of cytochrome c (cyt c) with non-deamidated (red, bottom) or deamidated (green, top) GII.4 Saga P dimer and 500 µM HBGA B ligand in 150 mM ammonium acetate solution at pH7. Signal intensity was normalized to the base peak in the spectra. The corrected HBGA B occupancy is shown as insets. (B) The corrected binding to non-deamidated and deamidated GII.4 Saga P dimer is shown for 500 µM and 250 µM Gb4.

Protein NMR experiments have recently demonstrated that MNV P dimers do not bind to HBGAs [22]. Moreover, MNV has altered dimerization properties and presents a large fraction of P monomers at neutral pH. While correction was not possible here due to insufficient quality of the reference protein signal (see Fig. S1), it is evident that similar glycan clustering is observed for P monomer and dimer. Glycan binding requires a dimeric protein, therefore, these interactions between MNV P monomers [26] and glycans must be unspecific in line with the study by Creutznacher et al. [22]. Additional support for largely unspecific interactions stems from a mutated hNoV GII.4 MI001 P dimer. Here, the supposed glycan binding pocket was mutated resulting in altered dimerization behaviour and hence P monomer and dimer signals. The corrected data shows equal clustering behaviour of HBGA B, A and GM3 on monomer and dimer (Fig. S2). Interestingly, Gb4 shows basically no binding after correction in this case. Noteworthy, correction for clustering based on P monomer results in no specific binding to the P dimer for the other three glycans as well. This suggests that cyt c is not a suitable reference and that glycan clustering does depend on biophysical properties of the proteins. Therefore, we compared several potential reference proteins, which are commercially available and exhibit distinct properties.

### 3.2 Glycan clustering to various reference proteins

The ratio, *R*, between the free to ligand-associated reference protein is used to eliminate non-specific clustering to the target protein. In total, eight different reference candidates, which differ in mass, size and structural composition, are examined displaying very different *R* values (Tab. 1). Exemplarily, the interaction of carbohydrates with the Saga P dimer and four reference proteins (cyt c, GFP, CA, and ADH) is shown (Fig. 2). The data illustrates similar binding patterns of HBGA B to Saga P dimers but vastly different glycan clustering to the reference proteins under identical measurement conditions. The dimeric ADH displays the strongest ligand clustering with ADH:ligand (1:1) becoming the base peak similar to the P dimer. CA and GFP show similar patterns with slightly less clustering, whereas cyt c presented a unique profile with lower clustering ratios and a non-Gaussian charge state distribution.

**Figure 2.**
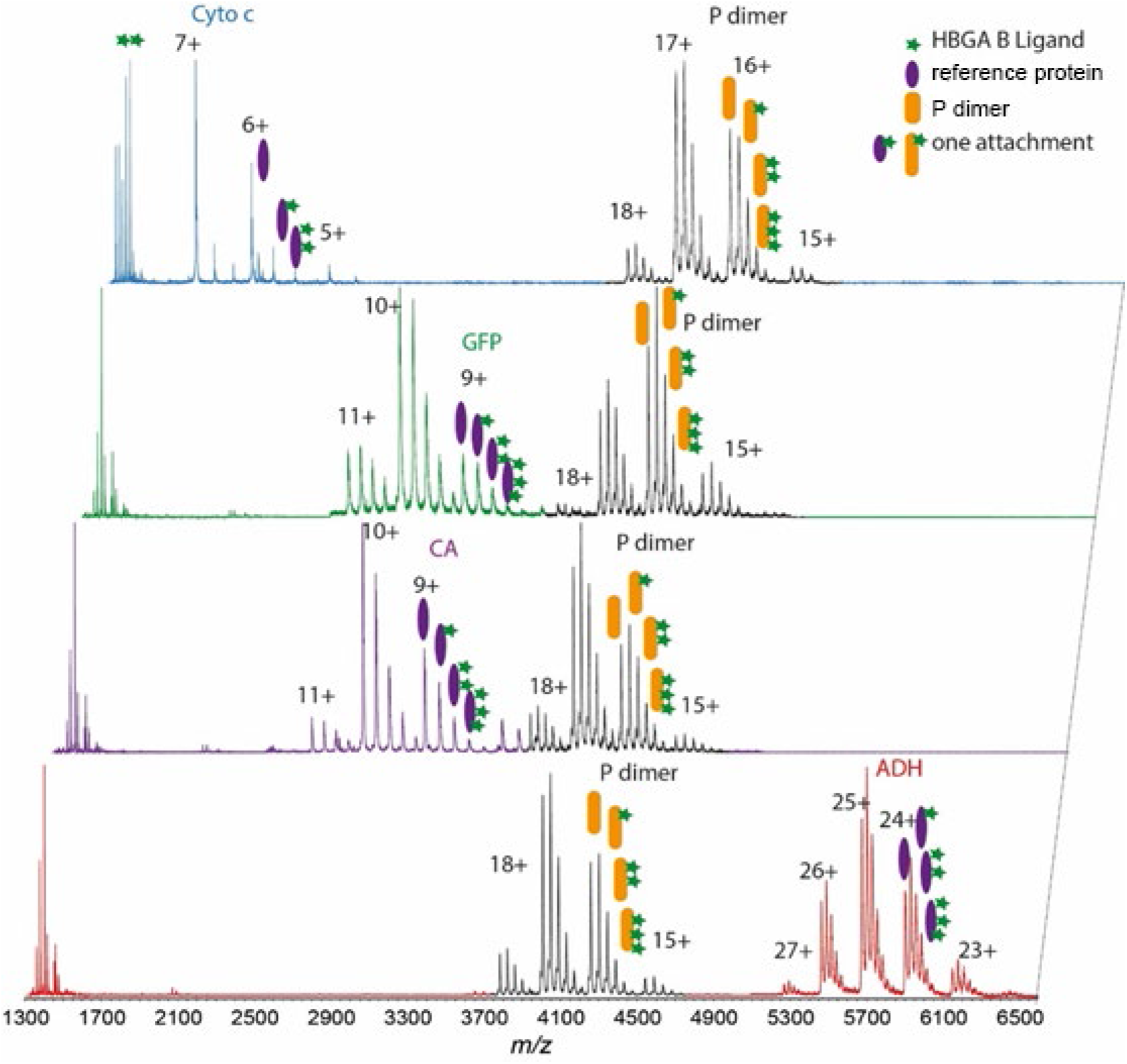
Native mass spectra of four selected reference proteins. 3 µM reference protein and 1 µM non-deamidated Saga P dimer (all protein concentrations based on the monomer) are measured with 500 µM HBGA B ligand in 150 mM ammonium acetate solution at pH 7. From top to bottom: cyt c (blue), GFP (green), CA (purple), ADH (red). Signal intensity was normalized to the protein base peak. P dimer mass range: 3800-5000 *m/z*, ADH mass range: 5200-6500 *m/z*, GFP mass range: 2700-3500 *m/z*, CA mass range: 2600-3700 *m/z*.

In line with the observed discrepancies between the reference proteins in *R* determination, the resulting K_D_ values for the same protein-ligand interaction vary strongly between a few 100 µM and 24 mM dependent on the choice of the reference protein. Notably, the K_D_ obtained with ADH as reference protein is in line to the expected value of 12 mM from NMR data [22] but also barely detectable. Differences in *R* and K_D_ are also consistent when titrating five glycan concentrations (Tab. S1). When the clustering ratio *R* is plotted over glycan concentration (Fig. 3), all reference proteins show the expected proportionality of *R* and glycan concentration for the three tested glycans but with different slopes. The plots clearly reveal strongest clustering to ADH in all conditions. On the other hand, the small-sized proteins (Myo, cyt c and Ubq, approx. 17 kDa, 13 kDa and 9 kDa) stick out with much lower clustering, the remaining proteins display intermediate *R* values. This could suggest an influence of molecular weight of the reference protein despite other reports [5,11]. While all plots fit to linear regression, the plots based on reciprocal *R* value reveal that clustering to Ubq is not linearly dependent on glycan concentration (Fig. 3, grey line). Therefore, Ubq is excluded from the following analysis.

**Figure 3.**
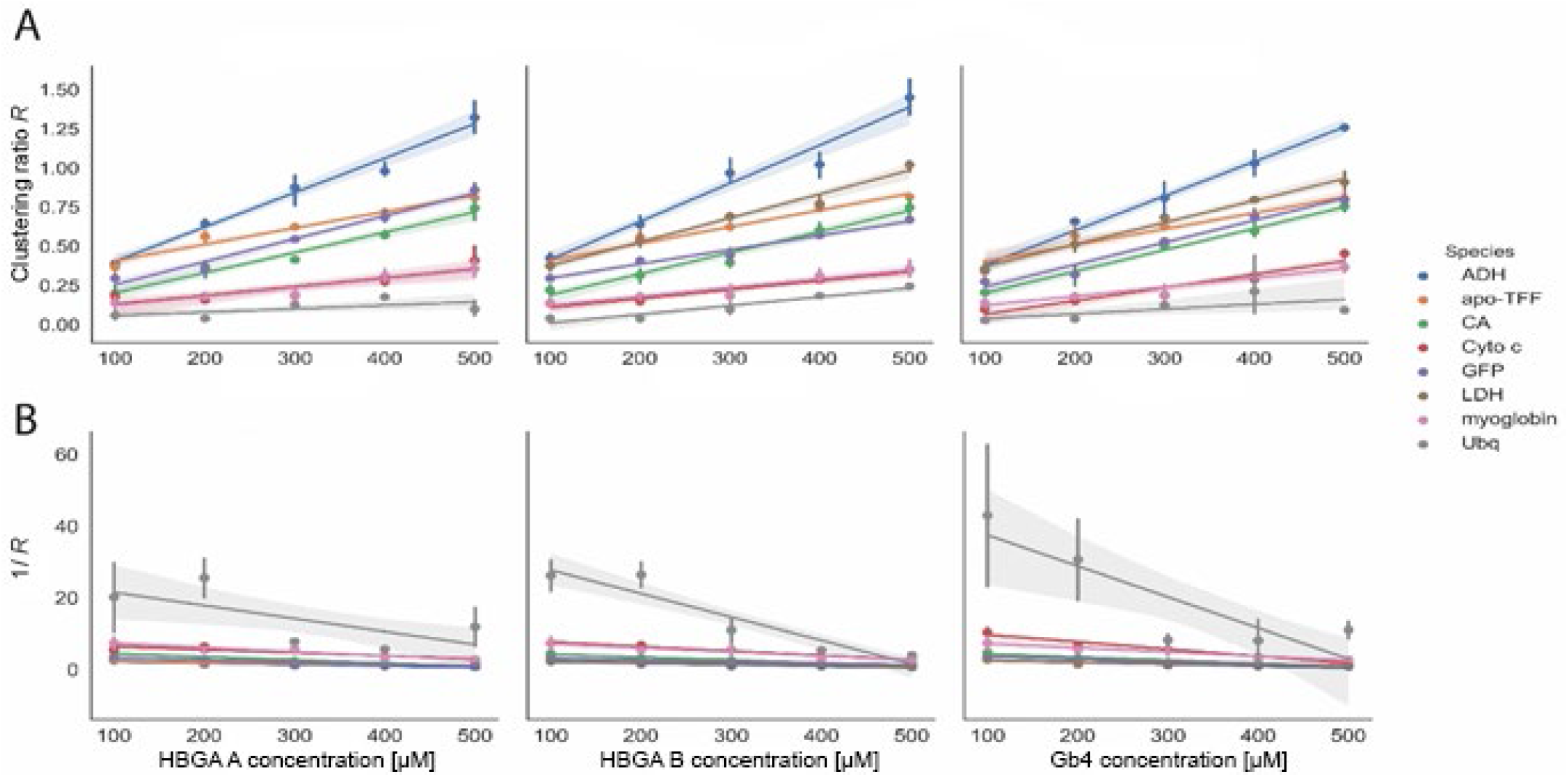
Correlating glycan concentrations to glycan clustering ratios. (A) *R* value calculated based on the peak-area of the reference proteins (ADH; CA, cyt c, GFP; Myo, Ubq, LDH, apo-TFF). (B) Corresponding reciprocal value. Titration experiments are performed with P dimers (1 µM, monomer) and reference proteins (3 µM, monomer) in 150 mM ammonium acetate solution pH 7 with ligand concentrations ranging from 100 µM to 500 µM for HBGA A, B and Gb4. The shaded areas represent the standard error of the slope for the linear fit.

### 3.3 Biophysical properties influence the glycan clustering

Ten different protein characteristics that could potentially affect glycan clustering are plotted against the *R* values of the remaining 7 reference protein candidates to dissect possible correlations: absolute amount of α-helix or β-sheet in kDa, number of total charged residues, total number of positively or negatively charged residues, isoelectric point (pI), *m/z*-values, molecular weight, solvent accessible surface area (SASA), collision cross section calculated from trajectory method (CCS TJM) (Fig. 4, Fig. S3 and S4). *R^2^* was used to assess the quality of the linear fits. The only parameter providing strong correlations at all ligand concentrations is the β-sheet amount of the reference proteins (Fig. 4), whereby β-sheet amount and glycan clustering grow proportionally. The absolute share of β-sheets in a protein is also related to the molecular weight. The larger a protein, the more amino acids can engage within β-sheets, which is the likely explanation for the apparent impact of protein size. Notably, the amount of α-helix shows no anti-proportional or any other correlation indicating that these contribute marginally to glycan clustering. Some of the parameters have *R^2^* values between 0.7 and 0.9, i.e. significantly lower than the β-sheet amount, however, a contribution to overall glycan clustering cannot be excluded for SASA, charged residues, molecular weight and *m/z*. CCS TJM and pI on the other hand are below 0.5 like α-helices and clearly do not contribute.

**Figure 4.**
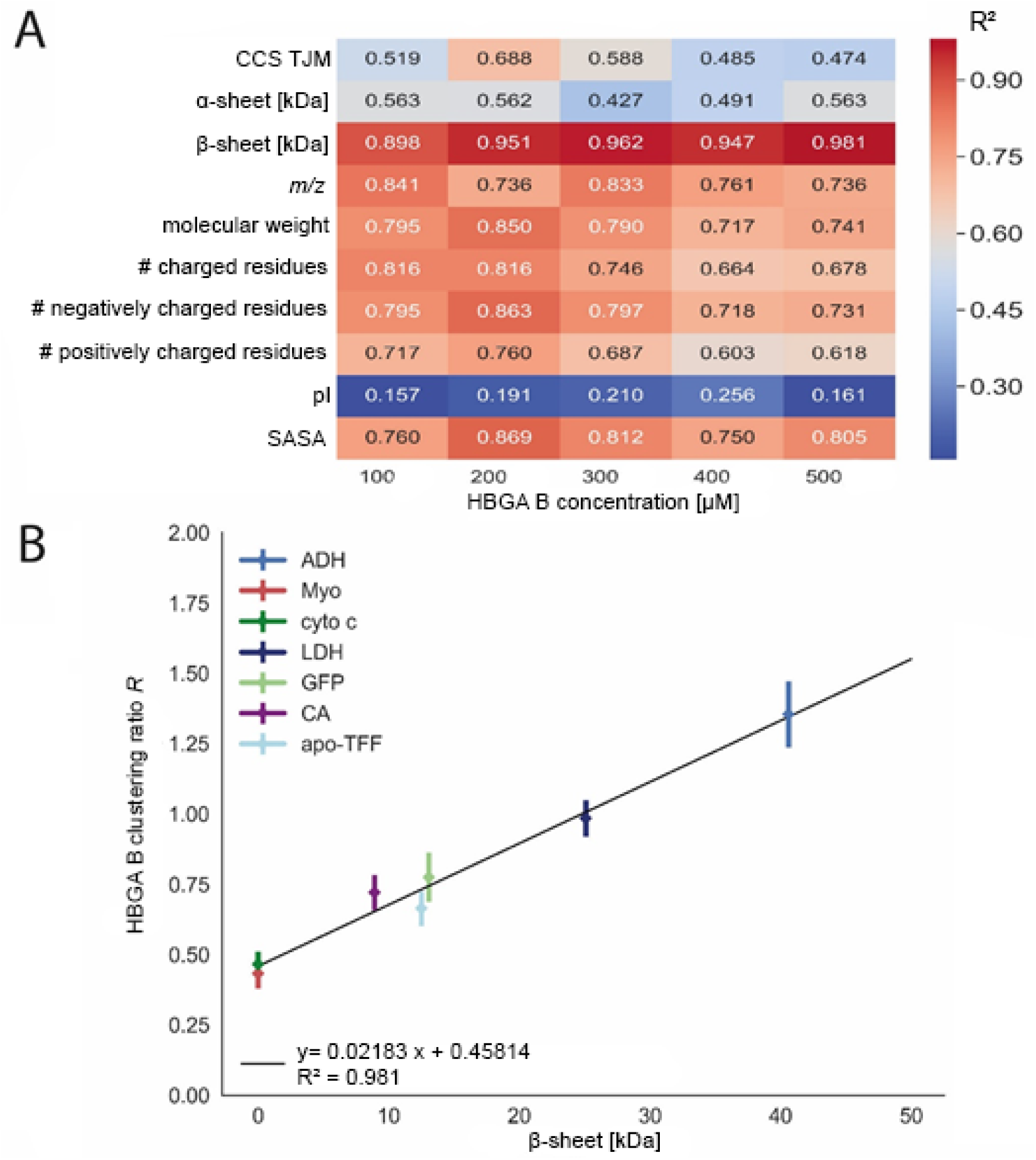
Correlation of clustering ratios *R* to different protein properties. (A) The heat-map of the *R^2^* value obtained from the linear regression at the indicated glycan concentration for the listed protein properties. (B) The correlation of β-sheet amount in kDa to unspecific glycan clustering ratios *R* at 500 µM HBGA B for seven reference proteins (ADH, CA, cyt c, GFP; Myo, LDH, apo-TFF). The black line represents the linear regression and the resulting equation is given.

### 3.4 Ion mobility on P dimer glycan interactions

Native ion mobility MS (IMMS) can reveal conformational changes in proteins, e.g. upon ligand binding, and has also been used to investigate P dimer glycan binding [27]. In IMMS, arrival time changes above the range of 3-5% are generally considered significant. A significant increase is not expected due to the binding of a small ligand like HBGA B in the absence of large conformational changes. This is also evident from HDX-MS and NMR [4]. In Figure 5, the arrival time distributions are shown for the non-deamidated GII.4 Saga P dimer together with its complexes with HGBA B (A) and Gb4 (B) of all detectable stoichiometries, and also respective single complexes of 1 P dimer : 0-3 HGBA B (C) and 1 P dimer : 0-3 Gb4 1 molecules (D). A small difference in the mean values and the peak widths can be observed among the panels A and B, and C and D in Figure 5, which is due to changes in environmental conditions, such as temperature, and slightly different tuning parameters used to optimize the signal. To ensure that no conformational changes were induced by the different conditions, the rotationally averaged collision cross section (CCS) for the pure P dimer was calculated: in A and C, the CCS was 3928 ± 14 Å^2^, while in B and D, the CCS was 3867 ± 12 Å^2^. The difference between these two CCS values is ∼1.6%, which is well below the significance threshold of the instrument, and is normal for experiments with large molecules. It can be seen that the arrival time distribution curves preserve their near-Gaussian, single-peak profile at all P dimer : ligand stoichiometries, suggesting no major conformational changes upon complex formation. Even though the arrival time distributions for all the complexes (Figure 5A and B) do not seem to follow an obvious trend, the difference between the arrival time distribution of the P dimer (black traces) and the sample with 500 µM of the respective ligands (green traces) suggest that at high ligand concentration, the relative concentration of the free P dimer is decreased. As it can be seen in the arrival time distributions of selected complexes (Figure 5C and D), with increasing number of bound ligands, the arrival time distribution curve shifts towards slightly higher values. The data clearly shows that the binding- and the non-binding ligand trigger the same behaviour, which we largely attribute to clustering.

**Figure 5.**
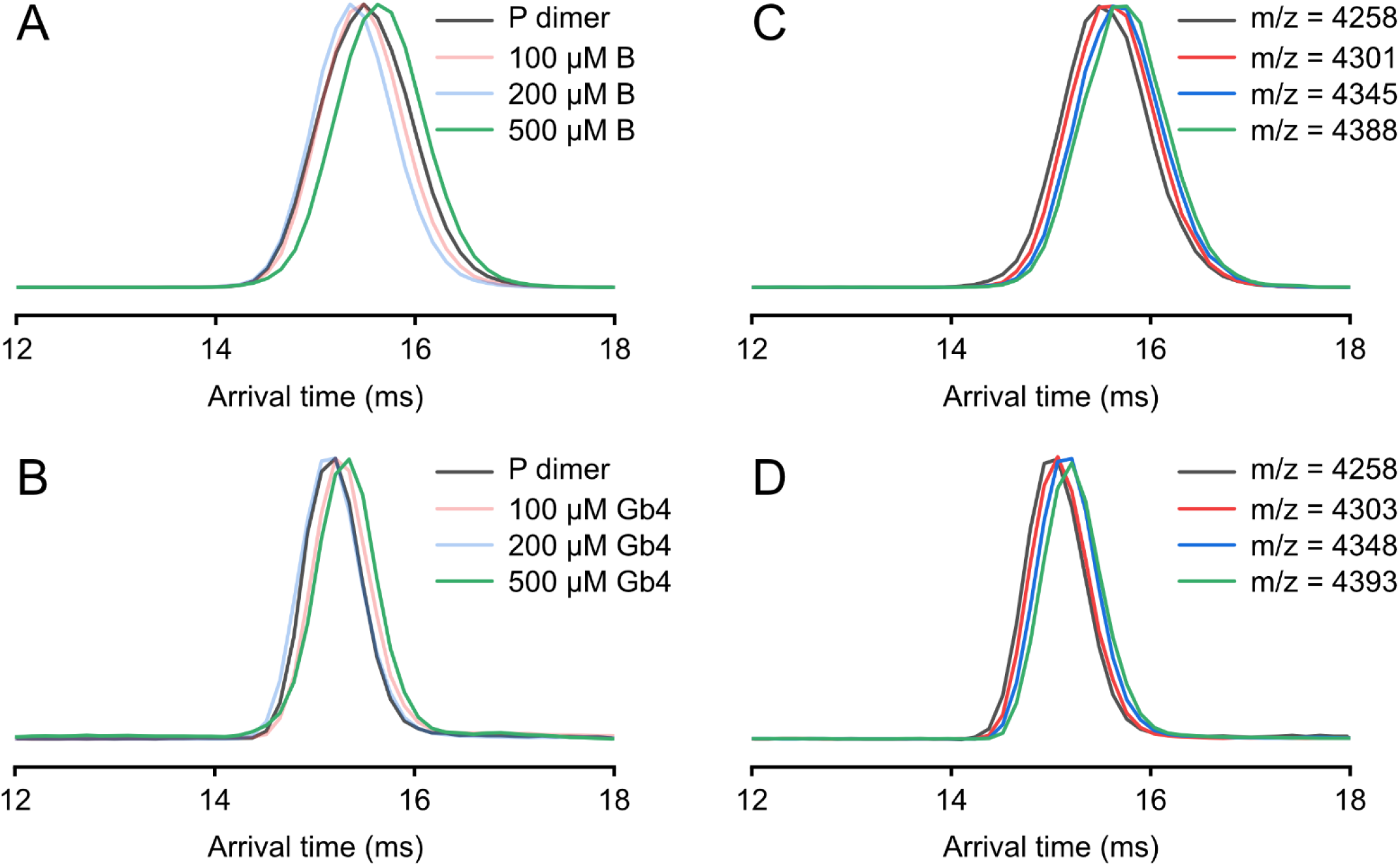
Glycan clustering to non-deamidated GII.4 P dimers in IMMS. Changes in the arrival time distribution upon increasing HBGA B (A) and Gb4 (B) concentration are depicted for the 16+ ion. Next to these are the extracted arrival time distributions for free P dimer and P dimer plus 1-3 ligands from a single measurement with HBGA B (C) and Gb4 (D) revealing an increasing trend with increasing number of ligands.

## 4. Discussion

Glycan clustering was observed previously in native MS experiments [5,9-11]. It occurs during the ionization process when solvent evaporates and the ions transfer to the gas phase. Due to excess of ligand, free glycans statistically end up in the same droplet and dry down onto the protein next to specific interacting glycans. The addition of a reference protein enables the determination of non-specific ligand clustering and hence correction to allow direct MS analysis of binding occupancy and K_D_. Therefore, this method has been widely used [12-14,28] to study protein-ligand interactions. Glycans pose a specific problem as the interactions are often of low affinity in the mM range requiring vast ligand excess to occupy binding sites.

Using deamidated GII.4 Saga P dimers as well as MNV and MI001 P monomers as negative controls for P dimer-glycan interaction, we reveal inherent problems with the direct MS approach employing reference proteins. This is further corroborated by complementary experiments using the non-binding all galactose glycan Gb4. The binding incompetent monomers show the same extent of glycan association as the respective P dimers. Of note, in these cases binding to the P dimers was also not expected due to mutations and MNV P dimers being unable to bind glycans at all [22]. Nevertheless for some glycans, MS yields even higher affinities for deamidated GII.4 Saga P dimers than for the wildtype, contradicting results from NMR and HDX-MS. HDX-MS showed increased flexibility in the deamidated P dimers suggesting that the structure affects glycan clustering. Here, we reexamine the degree of glycan attachment to different reference proteins to elucidate its origin and general suitability in native MS.

Therefore, we selected eight reference proteins differing in their properties. The clustering on ADH appears similar to the glycan distribution on the Saga P dimers while cyt c only presents a small amount of clustering. This further corroborates that glycan clustering is influenced by physicochemical properties of the protein. While most proteins show a linear correlation between clustering ratio and glycan concentration, Ubq behaves differently. In general, folded proteins are thought to ionize via the charged residue model (CRM). However, the non-linear glycan clustering behaviour of Ubq hints at ionization following another model as previously suggested by MD simulations. According to these simulations small proteins can also be ionized through the ion evaporation model (IEM) [29], which would explain the peculiar behaviour of Ubq and may also play a role for other small proteins. The cyt c clustering patterns have non-Gaussian character, which could be caused by ISD or varying ESI efficiencies. In contrast to Han *et al*., 2013 and 2018 [9,11], these results indicate unexpectedly different carbohydrate clustering propensity resulting in K_D_ spanning two orders of magnitude. The question arises, which biophysical characteristics cause this effect.

Previous work suggested that mass and size of reference proteins do not affect the correction procedure significantly [5,9]. Since most carbons in glycans carry hydroxyl groups, glycans have a much higher hydrogen bonding capacity than other small molecules, which could affect the interaction with the protein surface during ESI. Therefore, inspection of the structure of the selected reference proteins reveals distinct ratios of secondary structure elements. Various criteria are tested for correlation to the clustering ratio with the strongest correlation observed for β-sheet contents. We hypothesize that the net-like hydrogen bonding pattern in the β-sheets favors interaction with the glycans in contrast to α-helical structures. This could imply that intercalation occurs during the ionization process when glycans attach to the protein surface. Furthermore, β-sheets are more labile during the ESI process [30] which could facilitate glycan intercalation. Notably, reasonable correlation is also observed for molecular weight and SASA, which could be related to the higher probability of containing a significant amount of β-sheets with increasing size.

The direct MS approach with reference protein is therefore not well suited for studying low affinity glycan binding. It was originally developed for higher affinity interactions, where agreement in K_D_ to other methods was observed [5]. It also has proven invaluable for other ligand types [8,28,31]. IMMS demonstrated consistent results for HBGA B and Gb4 implying similar glycan clustering. No major changes in arrival time distributions are observed upon glycan addition in accordance with small ligands being added, and little structural changes observed in NMR and HDX-MS [4]. This is in contrast to a previous report [11], where the experimental and data acquisition parameters differed.

Overall, our results show that reference proteins with similar properties to the protein of interest should be used. Moreover, small proteins that could be affected by IEM upon ESI should be avoided. This explains some of the observed K_D_ discrepancies in literature. We have previously utilized cyt c, both small and mostly α-helical, therefore overestimating binding affinity [11,12]. Others used a small sized monoclonal antibody single chain fragment (scFv 26 kDa, [5,9,11]), which is mostly consiting of β-sheets. While the latter is much better suited, our data shows that protein dynamics as in the deamidated P dimer further influence the clustering [4,32].

The introduction of a non-binding ligand control is an additional assessment parameter to confirm the specific binding for low affinity ligands, which was also performed for glycan mimetics previously [13]. Comparing to stoichiometry information (Tab S2 and Fig. S5), the non-binder Gb4 expressed no specific binding in all listed glycan concentrations and HBGA B shows a single binding event at 500 µM concentration after ADH protein clustering correction. The calculated K_D_ (24 mM) is in accordance with the result from NMR (K_D_: HBGA B-tetrasaccharide type 1 12 mM, B trisaccharide 6.7 mM, fucose 22 mM) [4,22]. Hence, an appropriate reference candidate is crucial for the K_D_ determination of low affinity glycans.

## 5. Conclusion

In low affinity glycan binding studies, the K_D_ calculation is heavily dependent on the degree of glycan clustering on the reference protein. The protein structure and dynamics seem to influence the degree of glycan clustering heavily. Therefore, the quantification of direct binding affinity requires careful selection of a reference protein with similar β-sheet content and even structural dynamics to obtain accurate binding occupancy and affinities. The recent introduction of submicron emitters could be a way to reduce or circumvent the problem [33].

## Supporting information

complete supplement

## Supplementary Materials

All supplementary figures and tables are available online at www.mdpi.com/xxx/…

## Author Contributions

Conceptualization, C.U.; methodology, C.U., H.Y., J.L., G.P.S. and K.P.; software, H.Y.; investigation, H.Y., J.L., G.P.S.; resources, C.U., K.P., T.P., R.C., A.M. and S.T.; data curation, H.Y., J.L., G.P.S.; writing—C.U., J.L., H.Y., G.P.S.; writing—review and editing, all; supervision, C.U. and K.P.; project administration, C.U.; funding acquisition, C.U., T.P. and K.P. All authors have read and agreed to the published version of the manuscript.

## Institutional Review Board Statement

Not applicable.

## Informed Consent Statement

Not applicable.

## Data Availability Statement

All data is available from the corresponding author upon request.

