## Supplementary material for "Protein secondary structure affects glycan clustering in native mass spectrometry": complete supplement

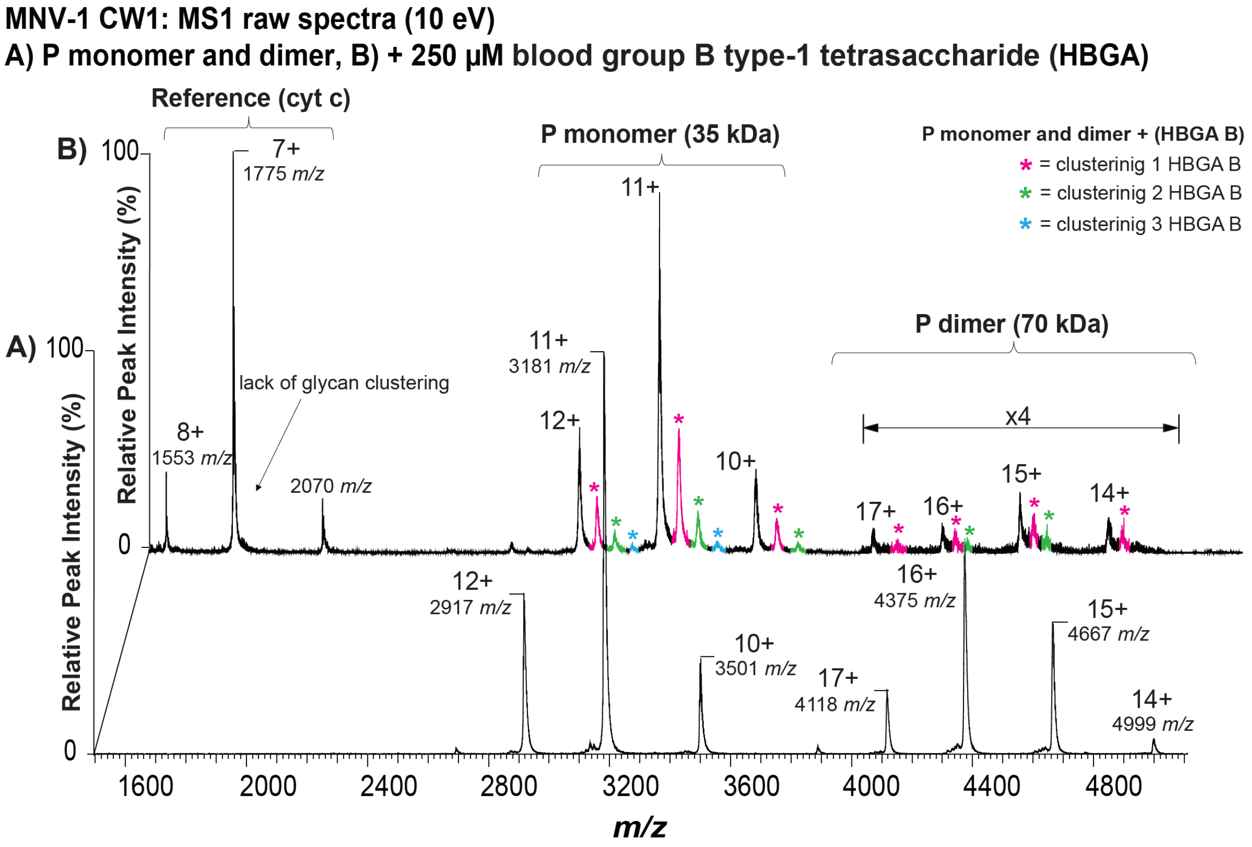


**Figure S1 glycan clustering on murine norovirus MNV-1 (CW1) P domain. A)** Native mass spectra of the P monomer and P dimer are shown, with a monomeric charge state distribution from 12+ to 10+ and a mass of 35 kDa and a dimeric charge state distribution of 17+ to 14+ with a mass of 70 kDa. **B)** Native mass spectra of cytochrome c (cyt c, 11 μM) with the P monomer (4 μM) and dimer (four times magnification) in presence of HBGA B clustering. HBGA B ligand concentration (250 μM) in 250 mM ammonium acetate solution at pH 7.0).

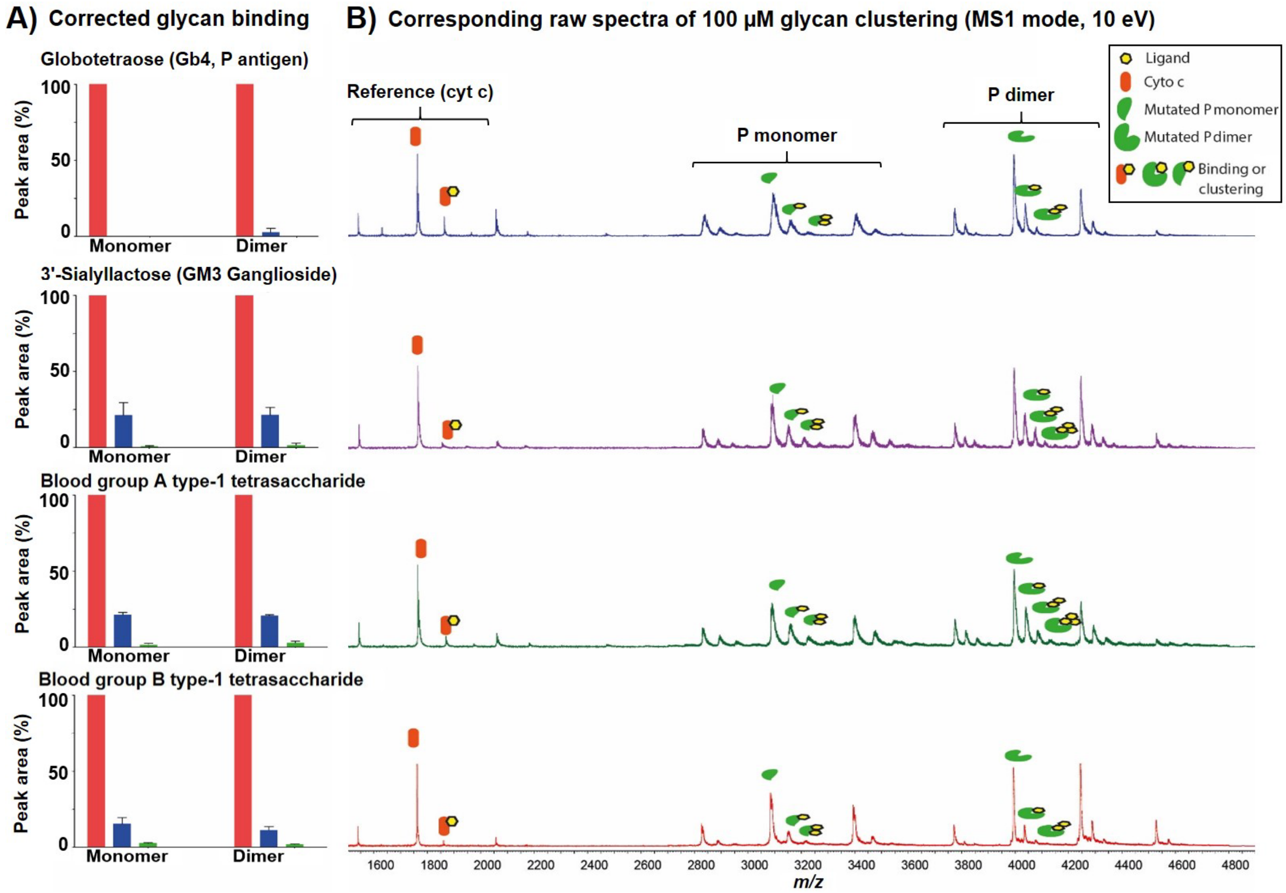


**Figure S2 glycan clustering on mutated hNoV GII.4 MI001 strain P domain. A)** The corrected binding of Gb4, GM3, blood group A and B type-1 tetrasaccharides (HBGA A, HBGA B) to mutated hNoV MI001 P dimer is shown, respectively in 100µM glycan concentration. **B)** Native mass spectra of cytochrome c (cyt c) with mutated hNoV MI001 P dimer in presence of different ligand Gb4 (dark blue), GM3 (purple), HBGA A (green), HBGA B (red) clustering (150 mM ammonium acetate solution at pH7). Signal intensity was normalized to the base peak in the spectra.

### Blood group B type-1 tetrasaccharide 300 µM

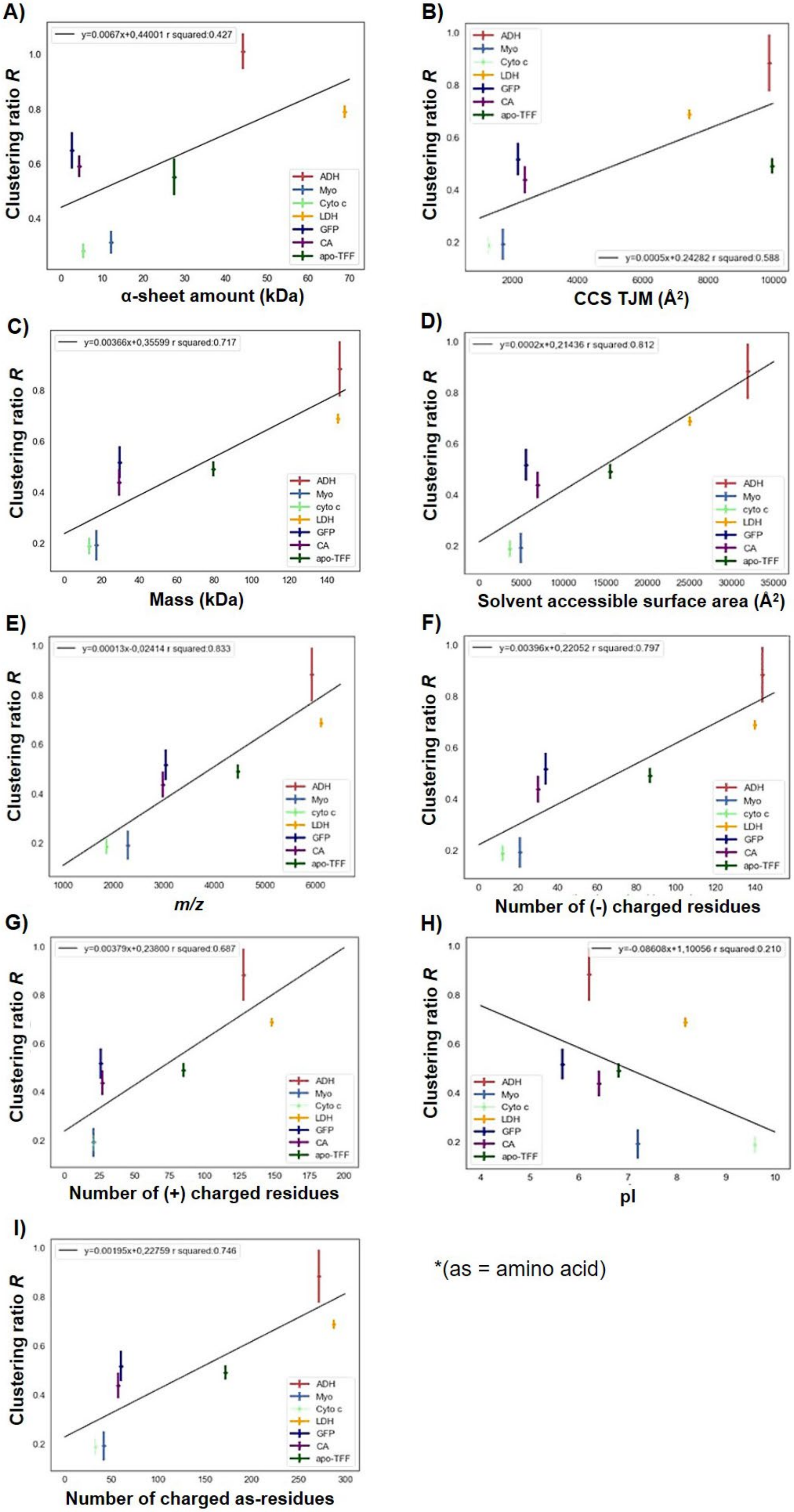


*(as = amino acid)

**Figure S3. Correlation of clustering ratios to multiple protein property parameters.** The correlation of different physiochemical parameters of protein (**A)** absolute share of $\alpha$-helix in a protein, **B)** CCS TJM, **C)** protein mass (kDa), **D)** solvent accessible area, **E)** mass to charge ratio ($m/z$), **F)** number of negative charged residues, **G)** number of positive charged residues, **H)** pI, **I)** number of charged residues) to unspecific glycan clustering ratios $R$ of seven reference proteins (ADH, CAII, cyt c, GFP, Myo. LDH, apo-TFF) at a blood group B type-1 tetrasaccharide (HBGA B) concentration of 300 µM was analysed. Native MS experiments were performed with a fixed concentration of P dimer (1 µM) and reference proteins (3 µM) at 150 mM ammonium acetate, pH 7. The reference protein structures and protein sequence information is derived from corresponding PDB files (see Table 1). The black line represents the linear regression.

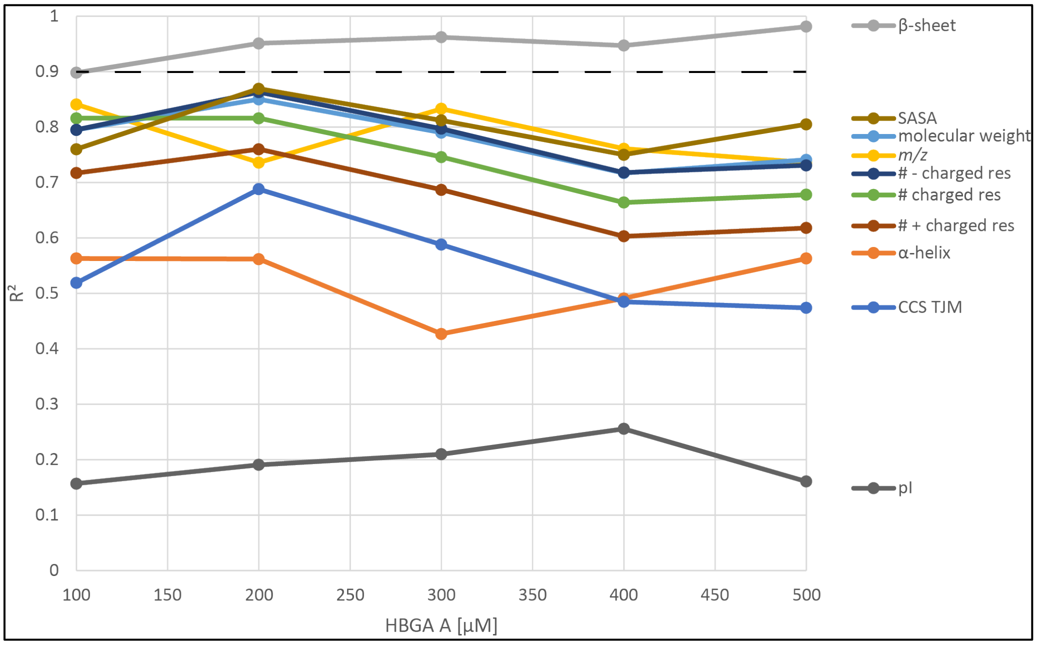


**Figure S4. Correlation plotted as $R^2$ of clustering ratios $R$ to different protein properties over employed HBGA A concentration.** $R^2$ of the different parameters (β-sheet amount in kDa, of $\alpha$-helix amount in kDa, charged residues, molecular weight of the protein, $m/z$) obtained for correlations at indicated HBGA A concentrations for the seven reference proteins. This is an alternative representation of the data displayed in Fig. 4A.

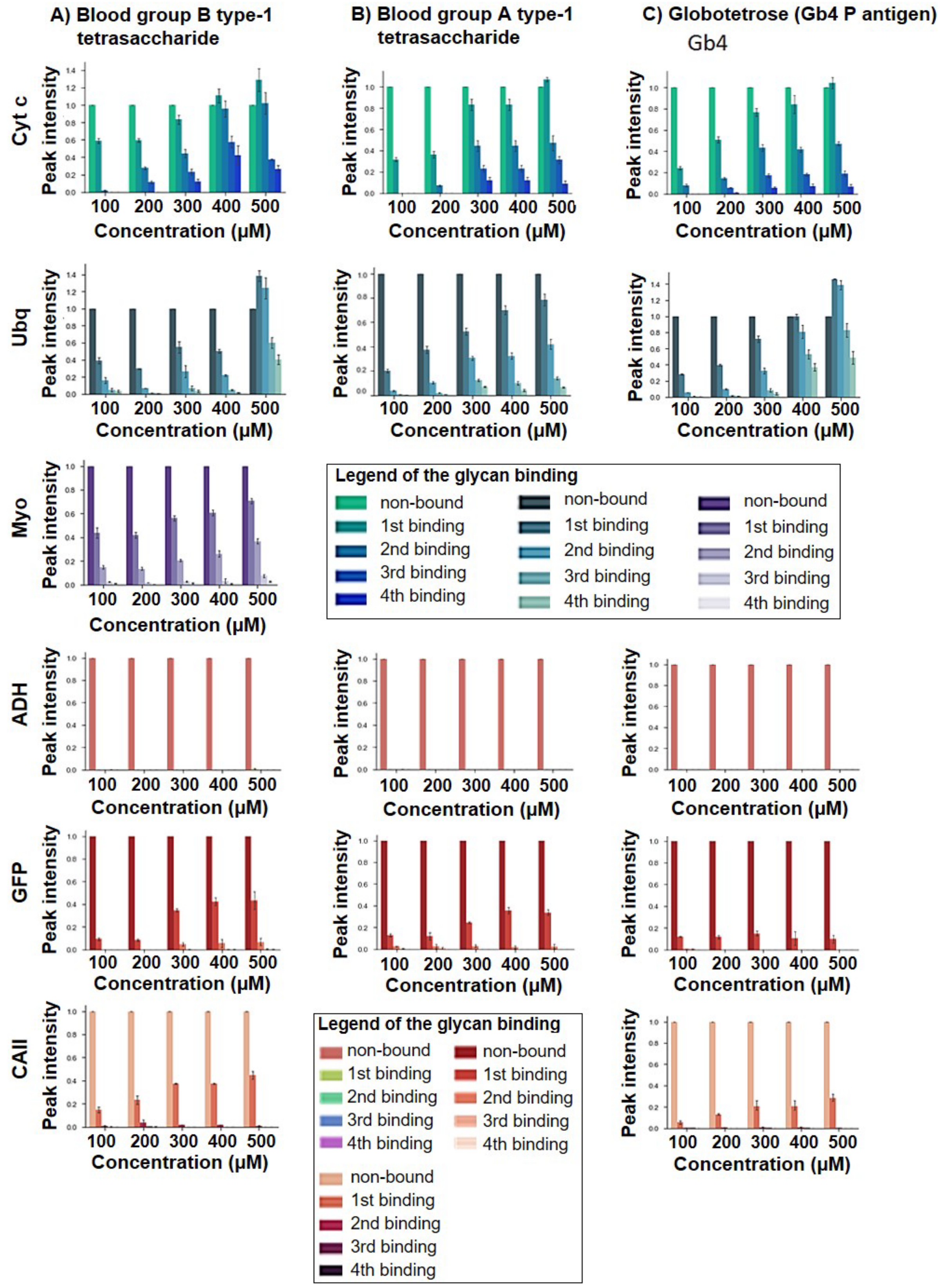


**Figure S5. Specific binding of different carbohydrates on the P dimer after correction of the unspecific glycan clustering.** Correction is based on the different reference proteins (glycans: blood group A and B type-1 tetrasaccharide (HBGA), Globotetrose (Gb4, P antigen) at 100 - 500 μM concentration; ref. proteins: cyt c, Ubq, Myo, ADH, GFP, CAII).

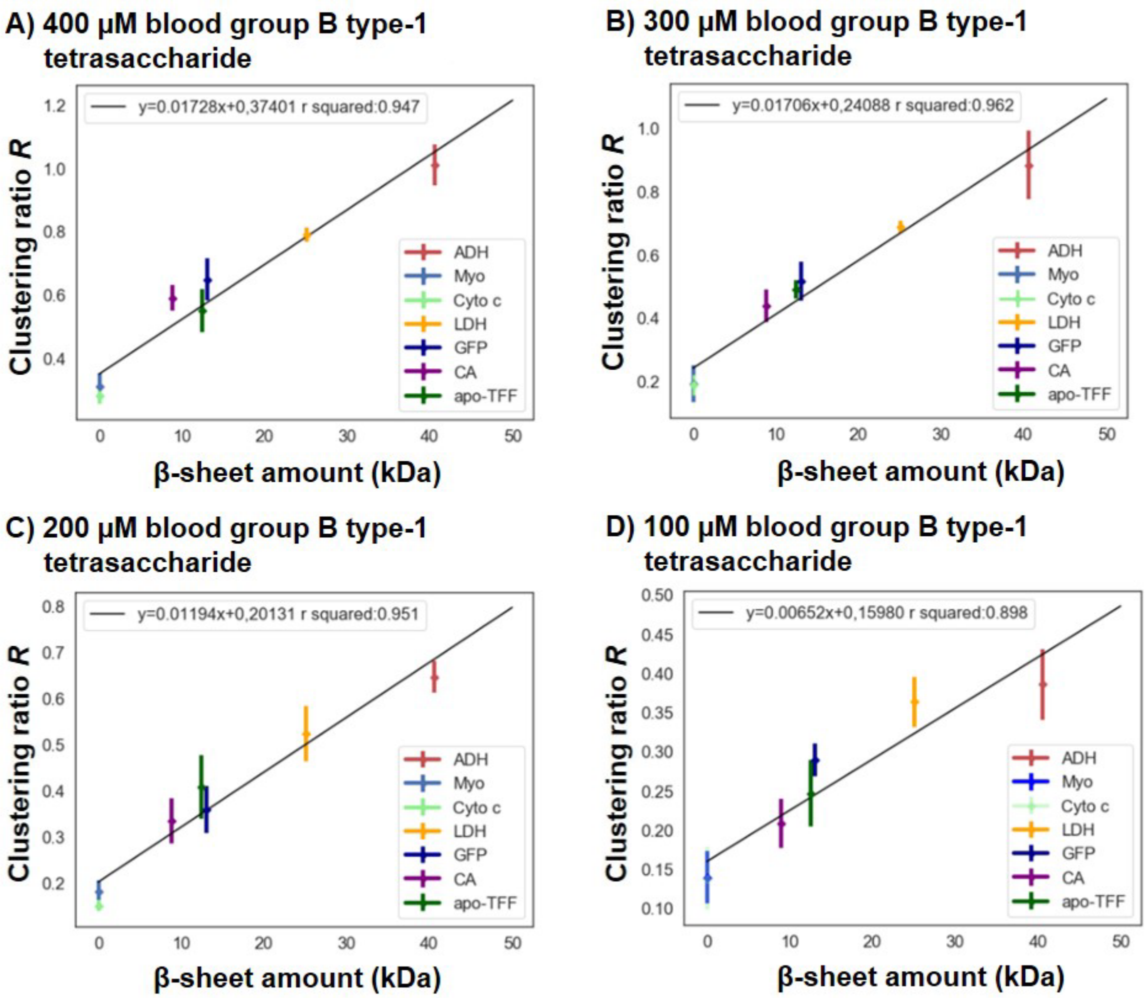


**Figure S6. The relation between the amount of beta-sheet in the reference protein and the unspecific glycan clustering ratio *R* at different ligand concentrations (400 μM (A), 300 μM (B), 200 μM (C) and 100 μM (D)).** The *R* value is calculated from the peak-area of the reference proteins (ADH, CA, cyto c, GFP, Myo, Ubq, LDH, apo-TFF). Native MS experiment were performed with a fixed concentration of P dimer (1 μM) and reference proteins (3 μM) at 150 mM ammonium acetate, pH7. The black line represents the linear regression between the unspecific clustering ratio of each reference protein and the corresponding mass of beta-sheets. The reference protein structures and protein sequence information is derived from each of corresponding defined PDB file.

**Table S1. Glycan clustering ratio of HBGA B interaction with P dimer analysed with native MS.** Data correction is based on the reference protein method.

| Ref. Protein | glycan clustering ratio | | | | | | | | | | | | | | |
|---|---|---|---|---|---|---|---|---|---|---|---|---|---|---|---|
| | 100 μM | | | 200 μM | | | 300 μM | | | 400 μM | | | 500 μM | | |
| | HBGA B | Gb4 | HBGA A | HBGA B | Gb4 | HBGA A | HBGA B | Gb4 | HBGA A | HBGA B | Gb4 | HBGA A | HBGA B | Gb4 | HBGA A |
| Myo | 0.14±0.03 | 0.14±0.02 | 0.13±0.03 | 0.18±0.02 | 0.17±0.03 | 0.16±0.03 | 0.19±0.06 | 0.20±0.03 | 0.22±0.03 | 0.31±0.04 | 0.30±0.01 | 0.29±0.04 | 0.36±0.05 | 0.37±0.06 | 0.34±0.03 |
| Cyt c | 0.14±0.01 | 0.10±0.01 | 0.18±0.03 | 0.15±0.01 | 0.15±0.00 | 0.16±0.02 | 0.19±0.03 | 0.19±0.03 | 0.19±0.03 | 0.29±0.02 | 0.29±0.04 | 0.27±0.02 | 0.35±0.03 | 0.45±0.02 | 0.41±0.09 |
| Ubq | 0.04±0.01 | 0.03±0.01 | 0.06±0.03 | 0.04±0.01 | 0.04±0.01 | 0.04±0.01 | 0.10±0.03 | 0.12±0.02 | 0.13±0.01 | 0.18±0.01 | 0.21±0.20 | 0.18±0.01 | 0.25±0.00 | 0.09±0.03 | 0.10±0.05 |
| GFP | 0.30±0.02 | 0.27±0.02 | 0.30±0.02 | 0.41±0.02 | 0.32±0.07 | 0.35±0.05 | 0.44±0.04 | 0.53±0.01 | 0.54±0.01 | 0.57±0.01 | 0.69±0.04 | 0.68±0.04 | 0.67±0.02 | 0.80±0.03 | 0.86±0.04 |
| CA | 0.22±0.03 | 0.21±0.01 | 0.20±0.05 | 0.32±0.05 | 0.32±0.07 | 0.37±0.01 | 0.40±0.04 | 0.50±0.00 | 0.42±0.02 | 0.60±0.05 | 0.60±0.05 | 0.57±0.02 | 0.75±0.07 | 0.75±0.03 | 0.74±0.07 |
| ADH | 0.42±0.03 | 0.34±0.04 | 0.39±0.02 | 0.64±0.06 | 0.66±0.03 | 0.65±0.02 | 0.97±0.08 | 0.81±0.11 | 0.88±0.10 | 1.02±0.08 | 1.03±0.08 | 0.98±0.05 | 1.45±0.11 | 1.26±0.03 | 1.32±0.10 |
| apo-TFF | 0.37±0.03 | 0.36±0.05 | 0.39±0.02 | 0.56±0.05 | 0.57±0.06 | 0.57±0.05 | 0.62±0.03 | 0.63±0.02 | 0.65±0.02 | 0.74±0.01 | 0.69±0.02 | 0.71±0.03 | 0.82±0.03 | 0.80±0.05 | 0.79±0.03 |
| LDH | 0.38±0.03 | 0.35±0.04 | n.a. | 0.54±0.05 | 0.51±0.08 | n.a. | 0.69±0.02 | 0.68±0.02 | n.a. | 0.77±0.05 | 0.80±0.01 | n.a. | 1.02±0.04 | 0.91±0.06 | n.a. |
| SAGA P dimer | 0.44±0.07 | 0.35±0.06 | 0.37±0.13 | 0.62±0.05 | 0.60±0.09 | 0.66±0.10 | 1.01±0.02 | 1.05±0.09 | 0.98±0.08 | 1.31±0.20 | 1.21±0.15 | 1.34±0.26 | 1.55±0.05 | 1.40±0.21 | 1.44±0.15 |

**Table S2. Bound number of carbohydrates on P dimer analysed with native MS.** Data correction is based on the reference protein method.

| Ref. Protein | Bound number of HBGA B on Saga P dimer | | | | | Bound number of HBGA A on Saga P dimer | | | | | Bound number of Gb4 on Saga P dimer | | | | |
|---|---|---|---|---|---|---|---|---|---|---|---|---|---|---|---|
| | 100 | 200 | 300 | 400 | 500 | 100 | 200 | 300 | 400 | 500 | 100 | 200 | 300 | 400 | 500 |
| Myo | 2 | 3 | 3 | 3 | 4 | 2 | 3 | 3 | 3 | 4 | 2 | 3 | 3 | 3 | 4 |
| Cyto c | 2 | 3 | 4 | 4 | 4 | 2 | 3 | 3 | 4 | 4 | 1 | 2 | 3 | 3 | 4 |
| Ubq | 2 | 3 | 4 | 4 | 4 | 2 | 3 | 4 | 4 | 4 | 2 | 3 | 4 | 4 | 4 |
| GFP | 1 | 1 | 2 | 2 | 2 | 1 | 1 | 1 | 1 | 1 | 2 | 2 | 2 | 2 | 2 |
| CA | 1 | 2 | 2 | 2 | 2 | 1 | 1 | 1 | 1 | 1 | 1 | 1 | 2 | 2 | 2 |
| ADH | 0 | 0 | 0 | 0 | 1 | 0 | 0 | 0 | 0 | 0 | 0 | 0 | 0 | 0 | 0 |
| apo-TFF | 1 | 1 | 1 | 1 | 2 | 0 | 0 | 1 | 1 | 1 | 0 | 0 | 1 | 1 | 1 |
| LDH | 0 | 0 | 0 | 0 | 1 | 0 | 0 | 0 | 0 | 1 | 0 | 0 | 0 | 0 | 1 |

**Table S3. Protein physiochemical properties used for plots**. Data is obtained from Protein Data Bank in Europe (https://www.ebi.ac.uk/pdbe-srv/pdbechem/)

| Ref. Protein | Analysed protein physiochemical feature | | | | | | | | |
|---|---|---|---|---|---|---|---|---|---|
| | The share of β-sheet | The share of α-helix | mass | CCS TJM | positive charge residues number | negative charge residues number | pI | charge residues number | solvent accessible area | *m/z* |
| ADH | 40.60 | 44.21 | 147.00 | 9876.81 | 128.00 | 144 | 6.21 | 272.00 | 31928.42 | 5933 |
| apo-TFF | 12.51 | 27.46 | 79.60 | 9959.74 | 85.00 | 87 | 6.81 | 172.00 | 15595.88 | 4469 |
| CA | 8.91 | 4.40 | 29.10 | 2378.38 | 27.00 | 30 | 6.41 | 57.00 | 6984.81 | 2985 |
| cyto c | 0.00 | 5.44 | 13.20 | 1286.00 | 21.00 | 12 | 9.59 | 33.00 | 3618.71 | 1864 |
| GFP | 13.08 | 2.61 | 29.60 | 2164.24 | 26.00 | 34 | 5.67 | 60.00 | 5608.96 | 3037 |
| LDH | 25.07 | 68.81 | 146.10 | 7421.70 | 148.00 | 140 | 8.17 | 288.00 | 25045.90 | 6120 |
| Myo | 0.00 | 12.14 | 17.00 | 1714.99 | 21.00 | 21 | 7.20 | 42.00 | 4980.50 | 2283 |
